# Persistent sensory processing and behavioral atypicalities in a mouse model of neonatal encephalopathy

**DOI:** 10.64898/2026.06.22.733663

**Authors:** Sakshi Narwekar, Mhasen Khalifa, Hannah M. Mulhern, Noah K. Simonds, Jennifer C. Burnsed, Adema Ribic

## Abstract

Neonatal hypoxia-ischemia (HI) injury is a major risk factor for lifelong cognitive impairments. Given its systemic impact, the neural mechanisms of impairments associated with HI injury remain unclear. In this study, we used a mouse model of neonatal HI injury to study its impact on goal-directed behavior and neural activity in adulthood using a head-fixed visual discrimination task. While neonatal HI injury did not impair discriminability or learning, it was associated with increased motor output in form of licking, faster reaction times and liberal decision bias, indicating an impulsive-like phenotype. These behavioral changes were accompanied by suppressed neuronal activity in the primary visual cortex (V1) and elevated cue-driven fluctuations in trial-to-trial firing variability in the prefrontal cortex (PFC), the latter of which was predictive of decision bias in HI mice. Our findings identify the long term impact of neonatal HI injury on goal-directed behavior, describe in detail the task-related patterns of neural activity in HI mice, and implicate abnormal neural variability in the PFC as a driver of impulsive-like behavior in adults that suffered neonatal HI injury.

## Introduction

One of the most common neurological conditions during the neonatal period is neonatal encephalopathy following a hypoxic-ischemic (HI) injury, with a prevalence of 3.5 per 1000 of live births in high income countries and up to 30 per 1000 live births in low-and middle-income countries (Acun et al., 2022; Gao & Jiang, 2024; Park et al., 2023). Neonatal HI is a systemic injury that disrupts blood flow and oxygenation, with up to 60% of surviving infants developing major neurological conditions, such as cerebral palsy, epilepsy, and behavioral and intellectual disabilities (Acun et al., 2022; Allen & Brandon, 2011). Structurally, 2 main patterns of brain injury have been described after neonatal HI (de Vries & Groenendaal, 2010): 1) the basal ganglia-thalamus pattern that correlates with severe motor impairments, such as cerebral palsy (Himmelmann et al., 2007), and 2) a more widespread “watershed” pattern of brain damage present in the white matter and the cortex that correlates with cognitive impairments (Steinman et al., 2009).

In animal models, HI injury results in a similar pattern of damage, with a gradient of basal ganglia-thalamus to cortical damage based on the timing of injury (Mallard & Vexler, 2015). The classical “Rice-Vannucci” model in postnatal mice and rats that involves unilateral carotid artery ligation followed by hypoxia (Rice et al., 1981), recapitulates both the basal ganglia-thalamus and the cortical damage, with volume loss in the hemisphere ipsilateral to the ligation (Brockmann et al., 2013; Kao et al., 2021). Mice with HI injury at postnatal day 7-10 (P7-10) show correlates of widespread neuronal activation in the hemisphere ipsilateral to the ligation, as well as motor deficits (Alexander et al., 2014; Du et al., 2026; Marlicz et al., 2024). While acute seizures are present immediately after the HI injury (Burnsed et al., 2019), neural activity measured through electroencephalography (EEG) is later dampened (Ranasinghe et al., 2015). Further, neurons in the prefrontal cortex (PFC), hippocampus, the visual (V1) and the retrosplenial cortex (RSC) show reduced firing rates long after HI injury (Brockmann et al., 2013; Domnick et al., 2015; Du et al., 2026; Failor et al., 2010).

In addition to motor and motor learning impairments (Du et al., 2026; Marlicz et al., 2024; Oorschot et al., 2007), mice and rats that suffered neonatal HI injury show sensory processing deficits (Alexander et al., 2014; Failor et al., 2010), as well as impairments in tasks that assess attention and impulsivity (Miguel et al., 2015, 2018). Previous studies hence suggest that structural and functional brain damage after neonatal and perinatal HI has a lasting impact on cognitive functions, but it is unclear how the reported neuronal function deficits relate to the reported behavioral phenotypes.

In this study, we used a well-established associative learning task (McCoy et al., 2026) to probe the behavior and neuronal activity during task performance in adult mice exposed to HI injury at P10. We found that neonatal HI mice display an impulsive-like phenotype, with liberal response bias and reduced reaction times to task cues. On a neuronal level, HI mice show impaired selectivity for task cues in the primary visual cortex (V1) and aberrant trial-to-trial firing rate variability in the prefrontal cortex (PFC), implicating sensory cue processing deficits in behavioral impairments after neonatal HI.

## Results

### Adult mice with neonatal HI injury show impulsive-like behavior in a visual discrimination task

Previous studies demonstrated that neonatal HI injury results in deficits in sensory processing and cognition (Failor et al., 2010; Miguel et al., 2015). To probe the neuronal basis of these deficits, we trained mice subjected to HI injury through left carotid ligation at postnatal day 10 (P10) on a head-fixed visual association task once they reached adulthood to facilitate electrophysiological recordings during task performance (Figure 1A) (McCoy et al., 2026). For this task, animals are implanted with stainless steel headposts at least 10 days before the onset of training, water restricted to 85% of their baseline weight after the recovery from implantation surgery and habituated to experimenters and the training environment before the onset of training. For the training, animals are head fixed over a conductive treadmill with their right eye that is contralateral to the HI injury centered on a computer monitor, as the mouse V1 receives stronger input from the contralateral eye (Ribic et al., 2019). The monitor displays the task cues and covers a significant portion of the right visual field (Figure 1B). The mice get their daily water allotment (1 ml) during task training through a spout positioned near their mouth that also records their licking responses (Hayar et al., 2006). 2 different cues are presented during the training: 120° and 60° oriented sinusoidal gratings at 100% contrast and 0.15 cpd (cycles per degree) spatial frequency, where the onset of 120° triggers the delivery of 7-10 µl of water through the spout. 60° orientation has no consequence. Animals were trained in 2 daily sessions, with each session consisting of 50 presentations of rewarded and non-rewarded cues each for a total of 100 trials. Each trial was classified using the licking responses into 4 groups: Hit (licking to 120°), False Alarm (licking to 60°), Miss (no licks to 120°) and Correct Rejection (no licks to 60°) (Figure 1C). Hits and False Alarms are used to quantify the behavioral performance (d’) according to the signal detection theory (SDT) (Green & Swets, 1966). For our task, we considered mice trained once they reached the d’ of 2 and maintained it for 3 consecutive sessions.

**Figure 1.**
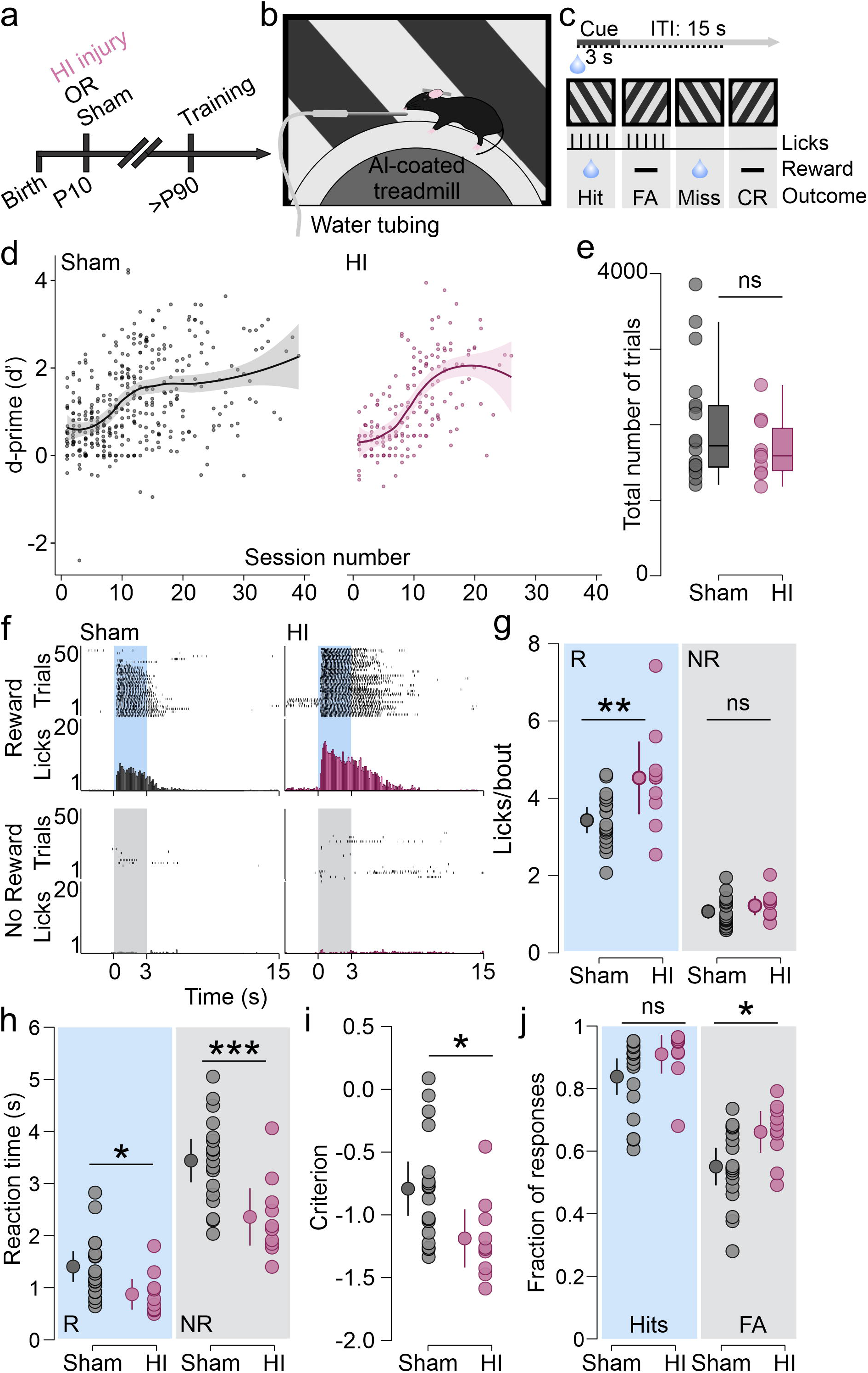
Neonatal HI injury results in impulsive-like behavior in adulthood. Schematics of **a)** experimental timeline, **b)** experimental setup, and **c)** task structure. **d)** Learning trajectories plotted as discriminability (d’) over time of sham (left) and HI (right) mice are not significantly different (LMM, sham vs HI β=0.04, p=0.493). **e)** Total number of trials to criterion is not different between sham and HI mice (Mann-Whitney t-test, p=0.38). **f)** Representative lick rasters of sham (left) and HI mice (right) to the rewarded cue (top) and non-rewarded cue (bottom). **g)** HI mice have a significantly increased number of licks following the rewarded cue (LMM, β=-0.54, p=0.005), but not following the non-rewarded cue (LMM, β=-0.07, p=0.31). **h)** Reaction times to both cues are significantly faster in HI mice (LMM: reward β=0.26, p=0.02; no reward β=0.52, p=0.002). **i)** Criterion values are significantly lower is HI mice (LMM, β=0.198, p=0.016). **j)** HI mice have a significantly increased fraction of False Alarms (LMM; Hits β=-0.036 p=0.097; False Alarms β=-0.055, p=0.016). d-j: data are represented as individual values from N=18 sham and 10 male and female HI mice. d: the line represents Loess fit and the shaded area 95% confidence intervals (CI). e: box plots represent median and interquartile range, with Tukey’s whiskers and individual values indicated. g-j: Estimated marginal means and the 95% CI are shown in addition to individual averaged values.

Both sham-operated (control) mice and mice with neonate HI injury performed as expected (McCoy et al., 2026), with low d’ values at the onset of training that gradually increased throughout the training (Figure 1D). We found no significant difference in the performance or learning trajectory between the two groups of mice [Figure 1E; linear mixed model (LMM), p_NeonatalHI_=0.49; N=18 sham and 10 HI mice of both sexes), and both groups required a similar number of trials to reach the training criterion (Mann-Whitney t test, p=0.38, U=71), demonstrating that neonatal HI injury has no impact on cue discriminability and leaning in a visual association task.

We next compared the licking responses between the two groups, as they represent the motor output in our task and provide information about the behavioral states (Matteucci et al., 2022). Mice of both groups licked similarly, with licks after the rewarded cue grouped closely after the cue onset, and very few licks present during and after the non-rewarded cue presentation once trained (Figure 1F). However, the HI mice had a significantly higher number of licks following the rewarded cue, despite similar duration of lick bouts (Figure 1F-G; LMM, Reward licks/bout p_NeonatalHI_=0.005; bout duration sham=8.28±0.26 s, HI=8.61±0.35 s, p_NeonatalHI_=0.45). Number of licks after the non-rewarded cue (during the False Alarm trials) was not significantly different between the two groups, despite longer bout duration in HI mice (Figure 1F-G; LMM, No Reward licks/bout p_NeonatalHI_=0.3; bout duration sham=6.78±0.22 s, HI=7.8±0.3 s, p_NeonatalHI_=0.009). The reaction times to both cues were significantly reduced in HI mice, indicating faster responding (Figure 1H; LMM, Reward p_NeonatalHI_=0.02, No Reward p_NeonatalHI_=0.001).

Faster reaction times are associated with a liberal decision bias (criterion or *c*) (Kloosterman et al., 2019), favoring responding over non-responding (Green & Swets, 1966). Indeed, HI mice had a significantly reduced *c* compared to sham mice, indicating a liberal decision bias (Figure 1I; LMM, p_NeonatalHI_=0.015). Liberal bias promotes responding and is associated with increased Hits and False Alarms, so we next compared the overall Hit and False Alarm rates between the two groups. While the Hits were not significantly different between sham and HI mice (LMM, p_NeonatalHI_=0.095), the rate of False Alarms was significantly higher in HI mice (Figure 1J; LMM, p_NeonatalHI_=0.015), indicating that HI mice commit more errors during the task. Trained sham and HI mice had similar motivation, calculated as trial-by-trial changes in lick latency during the rewarded trials (Berditchevskaia et al., 2016), with reaction times increasing as the session progressed (Supplementary Figure 1; LMM, reaction time p_trial_=0.04, p_neonatalHI_=0.15).

Altogether, our findings demonstrate that neonatal HI injury has a lasting impact on goal-directed behavior, with adult HI mice showing a liberal decision criterion, faster reaction times and increased error rates, suggesting an association between neonatal HI injury and impulsivity in adulthood (Edman et al., 1983).

### Processing of task cues is impaired in the V1 of HI mice

Neuronal activity in the primary visual cortex (V1) during cue presentation contributes to decision bias *c*, with heightened excitability promoting an increase in responding and a reduced *c* (liberal bias) (Jin & Glickfeld, 2019). We therefore tested if HI mice had altered neuronal activity in the V1 during task performance. To do that, we inserted a 16-channel linear probe spanning all cortical layers (NeuroNexus) into the craniotomy over the left V1 before task onset. After the probe settled, the spout was positioned into the place and the recordings of V1 activity from sham and HI mice were collected while the mice were performing the task (Figure 2A). Neuronal responses (single units) were isolated using template matching and threshold crossing and classified into narrow (putative fast-spiking (FS) interneurons) and wide (putative pyramidal neurons (RS) based on the shape of their waveform (Figure 2B; sham N=86 RS and 53 FS units, HI: N=227 RS units and 89 FS units) (Durand et al., 2016). After that, both types of waveforms were classified into 3 groups based on their cue-evoked responses: activated by task cues (“+”), suppressed by task cues (“-“) and non-responsive to task cues (“∼”, Figure 2C). We found that HI mice had a significantly reduced fraction of putative pyramidal neurons activated by the rewarded cue compared to sham animals, while the responsivity to the non-rewarded cue was intact (Figure 2C, left; 2-way RM ANOVA, cue×neonatal HI for Reward p=0.04, F_(2, 16)_=3.9; No Reward p=0.5, F=0.62; N=4 sham and 6 HI mice). Fractions of fast-spiking neurons activated and suppressed by task cues were comparable between sham and HI mice (Figure 2C, right; cue×neonatal HI for Reward p=0.11, F_(2, 16)_=2.5; No Reward p=0.5, F=0.73), indicating that the reduced representation of the rewarded cue in HI mice is specific to pyramidal neurons.

**Figure 2.**
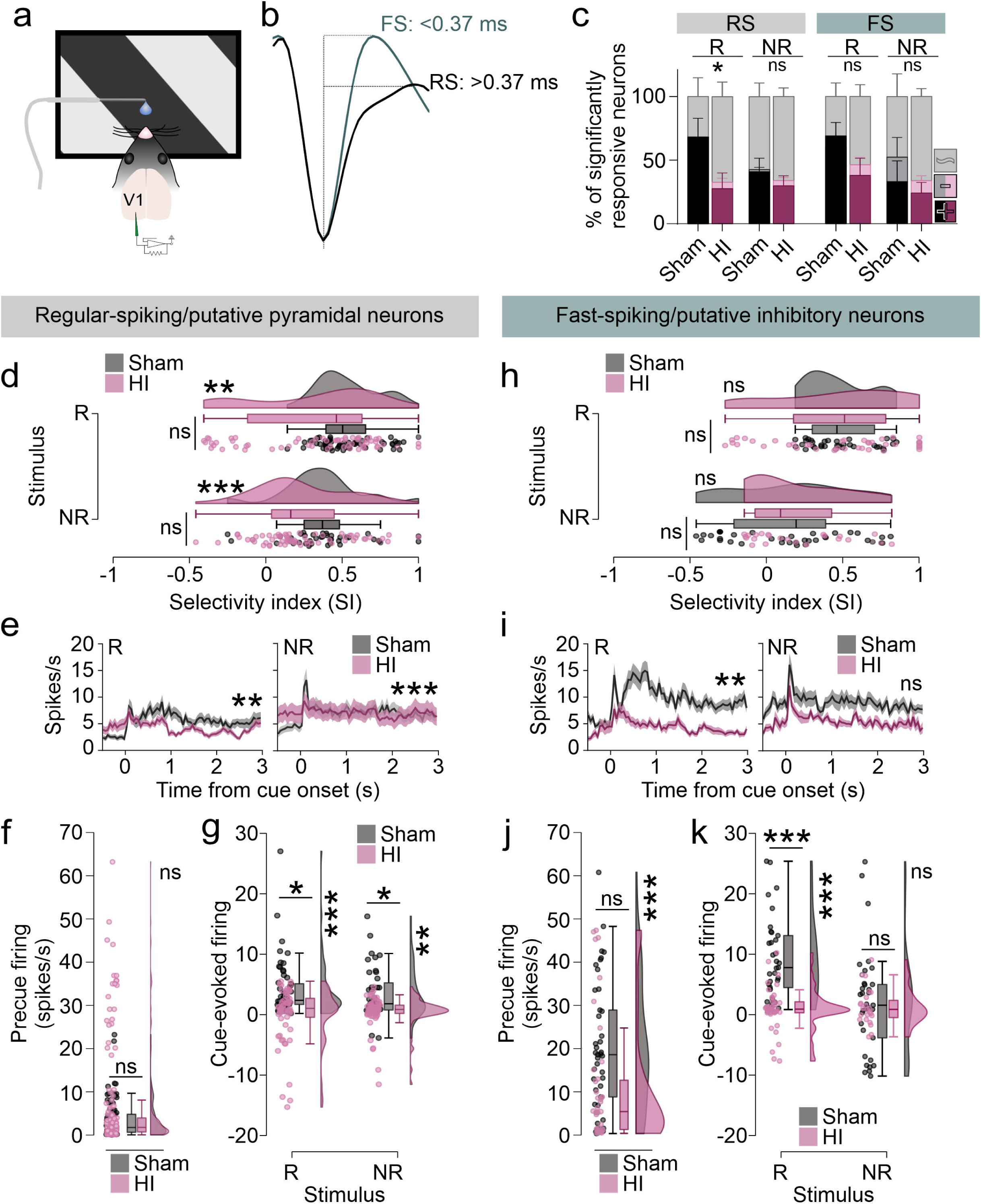
Processing of task cues is impaired in the V1 of trained HI mice. **a)**Schematics of experimental setup. **b)** Representative narrow (fast-spiking) and broad (regular-spiking) unit waveforms. **c)** Representation of task cues is reduced in regular-spiking (RS, grey bar) and intact in fast-spiking (FS, green bar) neurons in the V1 of HI mice (2-way RM ANOVA, RS: cue×neonatal HI for Reward p=0.04, F_(2, 16)_=3.9; No Reward p=0.5, F=0.62; FS: cue×neonatal HI for Reward p=0.11, F_(2, 16)_=2.5; No Reward p=0.5, F=0.73). **d)** Distribution of RS neuron selectivity indices for task cues is shifted towards lower values in the V1 of Hi mice (GLM; β_NeonatalHI_=0.08, p=0.26, β_Cue_=-0.063, p=0.01, β_NeonatalHI×Cue_=-0.014, p=0.57; KS test, p_Reward_=0.003, D=0.33; p_NoReward_=0.0007, D=0.4). **e)** PSTHs of units significantly responsive to or suppressed by task cues (from d). R=rewarded cue, NR=non-rewarded cue. Mann-Whitney t-test of peak response AUCs, p_Reward_=0.008, U=1227; p_NoReward_=3.61×10^-6^, U=587. **f)** Pre-cue firing of RS neurons is intact in HI mice (GLM, β_NeonatalHI_=0.104, p=0.79; KS test p= 0.4, D=0.15). **g)** Cue-evoked firing of RS units in HI mice is significantly blunted (GLM β_NeonatalHI_=1.63, p=0.035; β_Cue_=-0.05, p=0.84; β_NeonatalHI×Cue_=-0.189, p=0.47; Hommel post-hoc sham vs HI reward p=0.046, z=1.99, no reward p=0.046, z=2.05; KS test, p_Reward_=0.0005, D=0.37; p_NoReward_=0.0027, D=0.37). **h)** SI indices of putative fast-spiking interneurons are comparable between sham and HI mice (GLM, β_NeonatalHI_=0.029, p=0.75; β_Cue_=-0.115, p=0.02; β_NeonatalHI×Cue_=-0.03, p=0.51; KS test, p_Reward_=0.14, D=0.27; p_NoReward_=0.17, D=0.3). **i)** FS unit response to the rewarded cue is weaker in HI mice (Mann-Whitney t-test of AUC, p_Reward_=0.0085, U=600.5; p_NoReward_=0.46, U=543). **j)** Distribution of pre-cue firing rates of FS neurons in HI mice is shifted towards lower values (GLM, β_NeonatalHI_=0.2, p=0.49; KS test p= 0.0006, D=0.45). **k)** Responses to the rewarded cue are significantly lower in FS neurons of HI mice (GLM β_NeonatalHI_=2.46, p=0.021; β_Cue_=-1.33, p=0.14; β_NeonatalHI×Cue_=-1.85, p=0.05; Hommel post-hoc sham vs HI p_Reward_=2.4×10^-4^, p_NoReward_=0.71; KS test, p_Reward_=8.6×10^-8^, D=0.67; p_NoReward_=0.27, D=0.27). c: Data are represented as mean±SEM of 4 sham and 6 HI mice. e and i: Data are represented as mean±SEM of minimum 33 and maximum 64 neurons from 4 sham and 6 HI mice. d-k: box plots represent median and interquartile range, with Tukey’s whiskers and individual values indicated.

We next compared the firing properties of putative pyramidal and fast-spiking neurons in sham and HI mice that were classified as significantly activated or suppressed by task cues in more detail. We first compared the selectivity of both groups of neurons to task cues (rewarded and non-rewarded), quantified as the selectivity index (SI). SI values range from-1 to 1, with values closer to 1 indicating neurons highly selective for task cues. Average SI values of putative pyramidal neurons were not significantly different between sham and HI mice [Figure 2D; Generalized Linear Model (GLM); p_NeonatalHI_=0.26, p_Cue_=0.01, p_NeonatalHI×Cue_=0.57]. However, their distribution was significantly shifted towards lower values for both task cues in HI mice (Figure 2D, raincloud plots; KS test, p_Reward_=0.003, D=0.33; p_NoReward_=0.0007, D=0.4), indicating lower selectivity of V1 pyramidal neurons for task cues in HI mice on a population level. This was confirmed through comparison of peristimulus time histograms (PSTHs, Figure 2E), which revealed lower cue-evoked peak firing in HI mice for both task cues [Mann-Whitney t-test of area under the curve (AUC), p_Reward_=0.008, U=1227; p_NoReward_=3.61×10^-6^, U=587). Upon a more detailed comparison of pre-cue and cue-evoked firing, we found that HI and sham mice had comparable pre-cue firing rates (GLM, p_NeonatalHI_=0.79, KS test p= 0.4, D=0.15), but that HI mice had significantly reduced cue-evoked firing rates to both task cues and that their distribution was shifted to lower values as well (Figure 2F-G; GLM p_NeonatalHI_=0.035, p_Cue_=0.84, p_NeonatalHI×Cue_=0.47; KS test, p_Reward_=0.0005, D=0.37; p_NoReward_=0.0027, D=0.37).

SI indices of putative fast-spiking interneurons were comparable between sham and HI mice (Figure 2H; GLM, p_NeonatalHI_=0.75, p_Cue_=0.02, p_NeonatalHI×Cue_=0.51; KS test, p_Reward_=0.14, D=0.27; p_NoReward_=0.17, D=0.3). PSTHs revealed blunted firing to the rewarded cue in HI mice (Figure 2I, left) and a well differentiated peak after the non-rewarded cue presentation (Figure 2I, right; Mann-Whitney t-test of AUC, p_Reward_=0.0085, U=600.5; p_NoReward_=0.46, U=543). While the pre-cue firing rate of putative fast-spiking interneurons was not significantly different between sham and HI mice, their distribution was shifted towards lower values in HI mice (Figure 2J; GLM, p_NeonatalHI_=0.49; KS test p= 0.0006, D=0.45).

When we compared the cue-evoked firing rates, GLM identified a marginally significant interaction between the cue type and neonatal HI, as well as significant impact of HI injury, indicating a differential impact of neonatal injury on cue processing by fast-spiking interneurons (GLM p_NeonatalHI_=0.021, p_Cue_=0.14, p_NeonatalHI×Cue_=0.05). As expected from PSTHs, the firing evoked by the rewarded cue was significantly weaker in HI mice, and the distribution of firing rates was shifted towards lower values (Figure 2K; GLM, Hommel post-hoc p_Reward_=2.4×10^-4^; KS test, p_Reward_=8.6×10^-8^, D=0.67). Firing to the non-rewarded cue appeared intact in HI mice, both on the level of individual mice (Figure 2K, box plots; GLM, Hommel post-hoc p_NoReward_=0.71) and at the population level (Figure 2K, raincloud plots; KS test, p_NoReward_=0.27, D=0.27). Altogether, our findings demonstrate that neonatal HI injury impairs responsivity to cues during goal-directed behaviors in adult V1, confirming previous findings of reduced responsivity in the V1 after neonatal HI injury (Failor et al., 2010).

### Neonatal HI injury degrades the association between neural activity in the V1 and behavior

Neuronal activity in the V1 is associated with decision bias (*c*), such that optogenetic suppression and activation of V1 activity increases (more conservative) and decreases (more liberal) the values of *c*, respectively (Jin & Glickfeld, 2019). As HI mice had a significantly lower *c*, we tested the association between neuronal activity in the V1 and *c* in both groups of mice.

Cue-evoked firing of the regular spiking (putative pyramidal) neurons to either cue was not correlated with *c* in any of the mouse groups (Figure 3A-B; Pearson’s r: sham_Reward_=0.33, p=0.67; sham_NoReward_=-0.21, p=0.79; HI_Reward_=0.34, p=0.52; HI_NoReward_=0.31, p=0.55; N=4 sham and 6 HI mice). Rewarded cue-evoked firing of putative fast-spiking interneurons was also not associated with *c* (Figure 3C; Pearson’s r: sham_Reward_=-0.008, p=0.99; HI_Reward_=0.42, p=0.4). However, FS firing evoked by the non-rewarded cue was significantly correlated with c in the expected direction (Jin & Glickfeld, 2019), as well as predictive of *c* in sham mice only (Figure 3D; Pearson’s r: sham_NoReward_=-0.96, p=0.04; HI_NoReward_=0.63, p=0.18; linear regression R^2^ sham=0.92, HI=0.4). SI indices for task cues of putative pyramidal and fast-spiking neurons did not correlate with *c* in any of the mouse groups (Supplementary Figure 2; RS Pearson’s r: sham_Reward_=0.89, p=0.1; sham_NoReward_=-0.88, p=0.12; HI_Reward_=0.48, p=0.32; HI_NoReward_=0.6, p=0.2; FS Pearson’s r: sham_Reward_=0.29, p=0.7; sham_NoReward_=--0.91, p=0.09; HI_Reward_=-0.13, p=0.8; HI_NoReward_=0.14, p=0.79).

**Figure 3.**
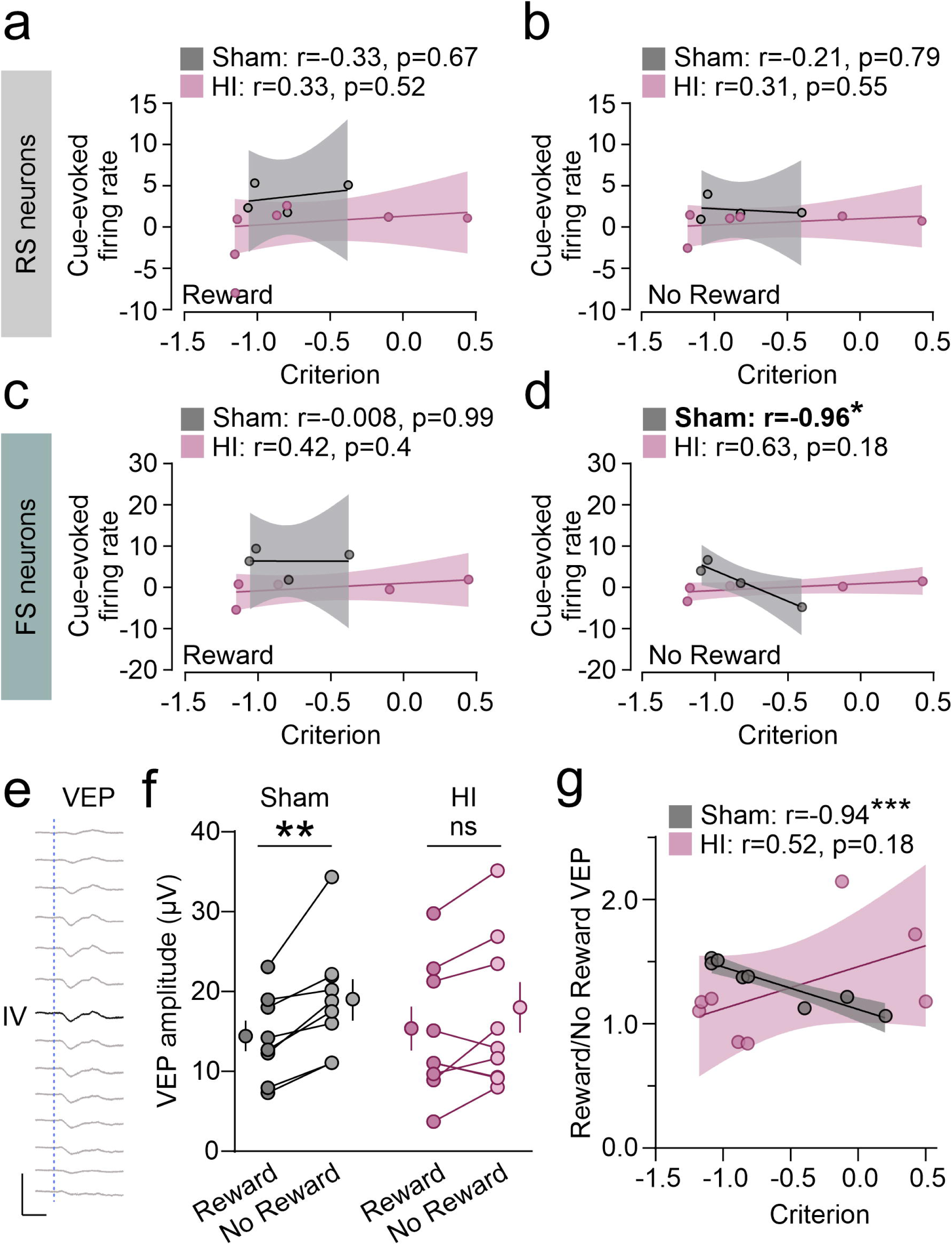
Neural activity in the V1 of HI mice is not correlated with decision bias. a-b) Cue-evoked firing of RS neurons to either cue is not correlated with criterion (*c*) in any of the mouse groups (Pearson’s r: sham_Reward_=0.33, p=0.67; sham_NoReward_=-0.21, p=0.79; HI_Reward_=0.34, p=0.52; HI_NoReward_=0.31, p=0.55; N=4 sham and 6 HI mice). **c)** Rewarded cue-evoked firing of FS neurons is not correlated with *c* (Pearson’s r: sham_Reward_=-0.008, p=0.99; HI_Reward_=0.42, p=0.4). **d)** Non-rewarded cue-evoked firing is significantly correlated with and predictive of *c* in sham mice only (Pearson’s r: sham_NoReward_=-0.96, p=0.04; HI_NoReward_=0.63, p=0.18; linear regression R^2^ sham=0.92, HI=0.4). **e)** Representative VEPs spanning cortical depth, with layer IV indicated in black. Scale bars: 100 ms and 100 µV. **f)** VEPs to the non-rewarded cue are significantly higher than VEPs to the rewarded cue in sham, but not in HI mice (LMM, β_NeonatalHI_=-0.01, p0.9, β_Cue_=1.77, p=5.4×10^-4^, β_NeonatalHI×Cue_=-0.5, p=0.23; Hommel post-hoc Reward vs No Reward: p_sham_=2.07×10^-4^, z=4.04; p_HI_=0.072, z=2.26). **g)** Ratio between VEPs to the non-rewarded and rewarded cue is significantly correlated with and predictive of *c* in sham mice, but not in HI mice (Pearson’s r sham p=0.0005, R^2^=0.88; HI p=0.18, R^2^=0.28; regression slope sham vs HI p=0.027). All points represent individual mice. a-d: N=4 sham and 6 HI mice. f-g: N=8 mice/group. f: Mean±SEM are indicated. a-f: 95% CI of linear regression line is indicated by shading.

To avoid spurious correlations as the number of sham mice we successfully isolated neurons from was low (N sham=4, HI=6), we analyzed cue-triggered local field potentials (LFPs) in this set of mice, as well as an additional set of mice from which we could not isolate single units. Neuronal spiking contributes to LFP (Buzsáki et al., 2012), which is known as the visually-evoked potential (VEP) in the V1 (Ribic et al., 2019). Given the relationship between the cue-evoked firing rates and *c* (Figure 3D), we hypothesized that VEPs would be also correlated with *c* in sham mice. We isolated VEPs in response to the rewarded and non-rewarded cues during task performance using linear silicone arrays and measured the amplitude of the negative peak (trough) in layer IV (the thalamorecipient layer; Figure 3E). While both sham and HI mice showed robust VEPs in response to task cues, the VEP in response to the non-rewarded cue was significantly higher than the VEP to the rewarded cue in sham, but not in HI mice (Figure 3F; LMM, p_NeonatalHI_=0.9, p_Cue_=5.4×10^-4^, p_NeonatalHI×Cue_=0.23; Hommel post-hoc Reward vs No Reward: p_sham_=2.07×10^-4^, p_HI_=0.072). Contrary to our initial hypothesis, the VEP amplitudes to the task cues were not correlated with *c* in sham mice (Pearson’s r Reward=0.002, p=0.99; No Reward=0.38, p=0.35). However, the ratio between them was significantly correlated with and predictive of *c* in sham mice, but not in HI mice (Pearson’s r sham p=0.0005, R^2^=0.88; HI p=0.18, R^2^=0.28; regression slope sham vs HI p=0.027, N=8 mice/group; Figure 3G). Our results demonstrate that decision bias *c* cannot be predicted from neural activity in adult HI mice, unlike from sham mice, indicating that neonatal HI injury degrades the association between neural activity in the V1 and behavior.

### Selectivity for task cues in the PFC is intact after neonatal HI injury

Neural signals propagate from the V1 to decision-making areas of the brain to influence behavior during tasks (Steinmetz et al., 2019). One such area is the prefrontal cortex (PFC), which also modulates cue selectivity in the V1 (Zhang et al., 2014). As neonatal HI injury disrupts the postnatal development of prefrontal networks (Brockmann et al., 2013), we next examined the long-term impact of HI injury on task-related neuronal activity in the PFC. We recorded the neuronal activity in the PFC (Figure 4A), encompassing the anterior cingulate and the prelimbic areas as previously described (McCoy et al., 2026). For analysis, we sorted the units into regular (RS) and fast-spiking (FS), as in Figure 2B.

**Figure 4.**
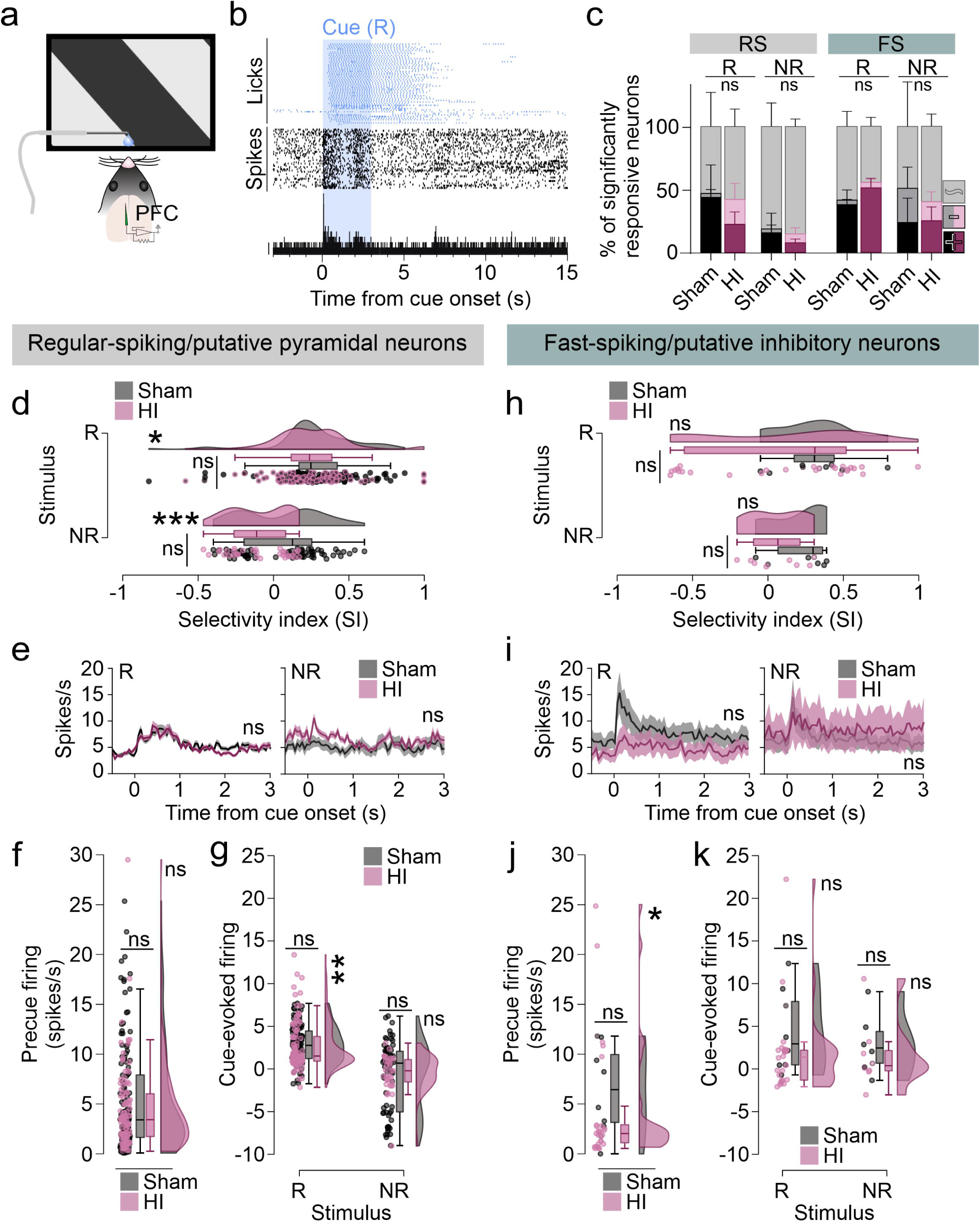
Processing of task cues is intact in the PFC of HI mice. **a)** Schematics of experimental setup. **b)** Representative PSTH (bottom) and rasters of unit firing (black) and licks (blue). **c)** Representation of task cues is intact in the PFC of HI mice (2-way RM ANOVA, RS: cue×neonatal HI for Reward p=0.37, F_(2, 18)_=1.02; No Reward p=0.86, F=0.13; FS: cue×neonatal HI for Reward p=0.63, F_(2, 16)_=0.47; No Reward p=0.83, F=0.19). **d)** Si indices of RS neurons are largely intact in HI mice, with only a minor shift in their distribution (GLM, β_NeonatalHI_=0.031, p=0.37, β_Cue_=-0.137, p=9.3×10^-4^, β_NeonatalHI×Cue_=0.014, p=0.64; KS test: p_Reward_=0.04, D=0.19; p_NoReward_=0.0003, D=0.44). **e)** PSTHs reveal similar firing patterns between RS neurons of sham and HI mice (Mann-Whitney t-test of AUC, p_Reward_=0.31, U= 4816; p_NoReward_=0.28, U=1047). **f)** Pre-cue firing of RS neurons is similar between sham and HI mice (GLM, β=0.085, p=0.6). **g)** Cue-evoked firing rates of RS neurons were similar between sham and HI mice, with the distribution shifted to lower values in HI mice (GLM, β_NeonatalHI_=-0.17, p=0.6; β_Cue_=-1.56, p=0.007; β_NeonatalHI×Cue_=-0.43, p=0.38; KS test, p_Reward_=0.0022, D=0.26; p_NoReward_=0.054, D=0.28). **h)** SI values of FS neurons were intact in HI mice (GLM, β_NeonatalHI_=0.61, p=0.48; β_Cue_=-0.015, p=0.84; β_NeonatalHI×Cue_=0.011, p=0.9), and their values were similarly distributed (raincloud plots; KS test: p_Reward_=0.45, D=0.35; p_NoReward_=0.27, D=0.5). **i)** PSTHs of FS neurons were also similar between sham and HI mice (Mann-Whitney t-test of AUC, p_Reward_=0.46, U= 120; p_NoReward_=0.78, U=40). **j)** Pre-cue firing of FS neurons was also intact in HI mice, despite a shift in the distribution towards lower values (GLM, β=1.08, p=0.34; KS test p=0.03, D=0.58). **k)** Cue-evoked firing of FS neurons was similar between sham and HI mice (GLM, β_NeonatalHI_=0.74, p=0.42; β_Cue_=-0.26, p=0.7; β_NeonatalHI×Cue_=-0.71, p=0.51; KS test, p_Reward_=0. 22, D=0.42; p_NoReward_=0.46, D=0.42). c: Data are represented as mean±SEM of 5 sham and 6 HI mice for RS, and 4 sham and 6 HI mice for FS. e and i: Data are represented as mean±SEM of minimum 6 and maximum 104 neurons from 4 (FS)/5 (RS) sham and 6 HI mice. d-k: box plots represent median and interquartile range, with Tukey’s whiskers and individual values indicated.

Neuronal spiking of both unit types was strongly driven by task cues and preceded the licking responses (Figure 4B). Cue representation, calculated as the fraction of neurons (per mouse) significantly driven or suppressed by task cues, was not impacted by neonatal HI in any of the two groups of isolated neurons (Figure 4C; 2-way RM ANOVA, RS: cue×neonatal HI for Reward p=0.37, F_(2, 18)_=1.02; No Reward p=0.86, F=0.13; FS: cue×neonatal HI for Reward p=0.63, F_(2, 16)_=0.47; No Reward p=0.83, F=0.19; N=5 sham for RS and 4 for FS, and 6 HI mice).

Selectivity indices of putative pyramidal neurons significantly modulated by task cues were not significantly different between the two groups (Figure 4D; GLM, p_NeonatalHI_=0.37, p_Cue_=9.3×10^-4^, p_NeonatalHI×Cue_=0.64). However, the distribution of putative pyramidal neuron SIs in HI mice was shifted towards lower values (Figure 4D, raincloud plots; KS test: p_Reward_=0.04, D=0.19; p_NoReward_=0.0003, D=0.44). Unlike in the V1 (Figure 3E and I), PSTHs revealed similar firing patterns between sham and HI mice (Figure 4E; Mann-Whitney t-test of AUC, p_Reward_=0.31, U= 4816; p_NoReward_=0.28, U=1047). Analysis of pre-cue and cue-evoked firing further confirmed no significant differences in the firing of putative pyramidal neurons in the PFC of HI mice, except for the distribution shift towards lower values in HI mice (Figure 4F-G; GLM, pre-cue firing rate sham vs HI p=0.6; cue-evoked firing rate p_NeonatalHI_=0.6, p_Cue_=0.007, p_NeonatalHI×Cue_=0.38; raincloud plots, KS test, p_Reward_=0.0022, D=0.26; p_NoReward_=0.054, D=0.28).

Selectivity of putative fast-spiking interneurons was similar between sham and HI mice (Figure 4H; GLM, p_NeonatalHI_=0.48, p_Cue_=0.84, p_NeonatalHI×Cue_=0.9), and their values were similarly distributed (Figure 4H, raincloud plots; KS test: p_Reward_=0.45, D=0.35; p_NoReward_=0.27, D=0.5). PSTHs were also similar (Figure 4I; Mann-Whitney t-test of AUC, p_Reward_=0.46, U= 120; p_NoReward_=0.78, U=40), and detailed analysis of firing revealed that while the pre-cue firing rates were overall shifted towards lower values in HI mice (Figure 4J; GLM, p=0.34; KS test p=0.03, D=0.58), both cues evoked similar responses in putative fast-spiking interneurons of sham and HI mice (Figure 4K; GLM, p_NeonatalHI_=0.42, p_Cue_=0.7, p_NeonatalHI×Cue_=0.51; KS test, p_Reward_=0. 22, D=0.42; p_NoReward_=0.46, D=0.42).

Compared to the V1, neuronal firing in the PFC appeared less impacted by the neonatal HI injury (Figures 2 and 4). Indeed, when we measured the thickness of the PFC in sham and HI mice we found no significant differences, despite damage to the hippocampus on the side ipsilateral to the injury in HI mice (Supplementary Figure 3). Cortical thickness ipsilateral to the injury side was not correlated with either d’ or *c* (Pearson’s r, d’=0.011, p=0.99; *c*=-0.171, p=0.53; N=12 sham and 10 HI mice). Altogether our results indicate that neonatal HI injury transiently impacts the neuronal activity in the PFC (Brockmann et al., 2013).

### Neonatal HI injury impairs the temporal dynamics of neuronal firing rate variability in the PFC

Neuronal firing in the PFC mediates performance and reaction times in associative tasks (Kim et al., 2021; McCoy et al., 2026), both of which are significantly different between HI mice and sham controls (Figure 1H-I). As selectivity for and the representation of task cues was intact in the PFC of HI mice, we next tested if neonatal HI injury impacts other aspects of neuronal firing, such as firing rate variability, recently highlighted as a potential biomarker of neuropsychiatric conditions (Tsikonofilos et al., 2025; Waschke et al., 2021). Neuronal trial-to-trial firing rate variability, calculated as the Fano factor (the ratio between trial-to-trial variance and mean spike count), is quenched by cue presentation in the V1 (Churchland et al., 2010) and in the PFC after task training (Figure 5A) (Qi & Constantinidis, 2012; Wu et al., 2017). We calculated Fano factor in 100 ms time bins 0.5 s before (baseline) and 1.5 s after the presentation of the rewarded cue for all isolated units (183 units from 5 sham and 294 units from 6 HI mice) and compared its values between sham and HI mice (Figure 5B), as well as temporal dynamics (Figure 5C) using two-sided cumulative sum (CUMSUM) and PELT (Pruned Exact Linear Time) changepoint detection. Both groups of mice showed a comparable suppression of trial-to-trial firing rate variability (Fano factor; ∼12–13%) following the rewarded cue presentation (Figure 5B and C; GLM, p_NeonatalHI_=0.79, p_Cue_=0.004, p_NeonatalHI×Cue_=0.55; N=5 sham and 6 HI mice). Two-sided CUSUM further revealed that the Fano factor of prefrontal neurons in sham mice returned to ±1 SD of baseline by 0.35 s post-cue and remained within those values. Fano factor of prefrontal neurons in HI mice recovered from the post-cue trough at a similar rate, but instead of settling at ±1 SD of baseline like in sham mice, it went past it and reached ∼22% above the baseline level between 0.65 and 1.25 s (Figure 5C). PELT changepoint detection identified a structural break in the HI Fano factor at 0.55 s, marking a transition from a below-baseline segment (mean z=-1.14) to an above-baseline segment (mean z=+0.63). No such break was detected in the Fano factor values of sham mice (Figure 5C).

**Figure 5.**
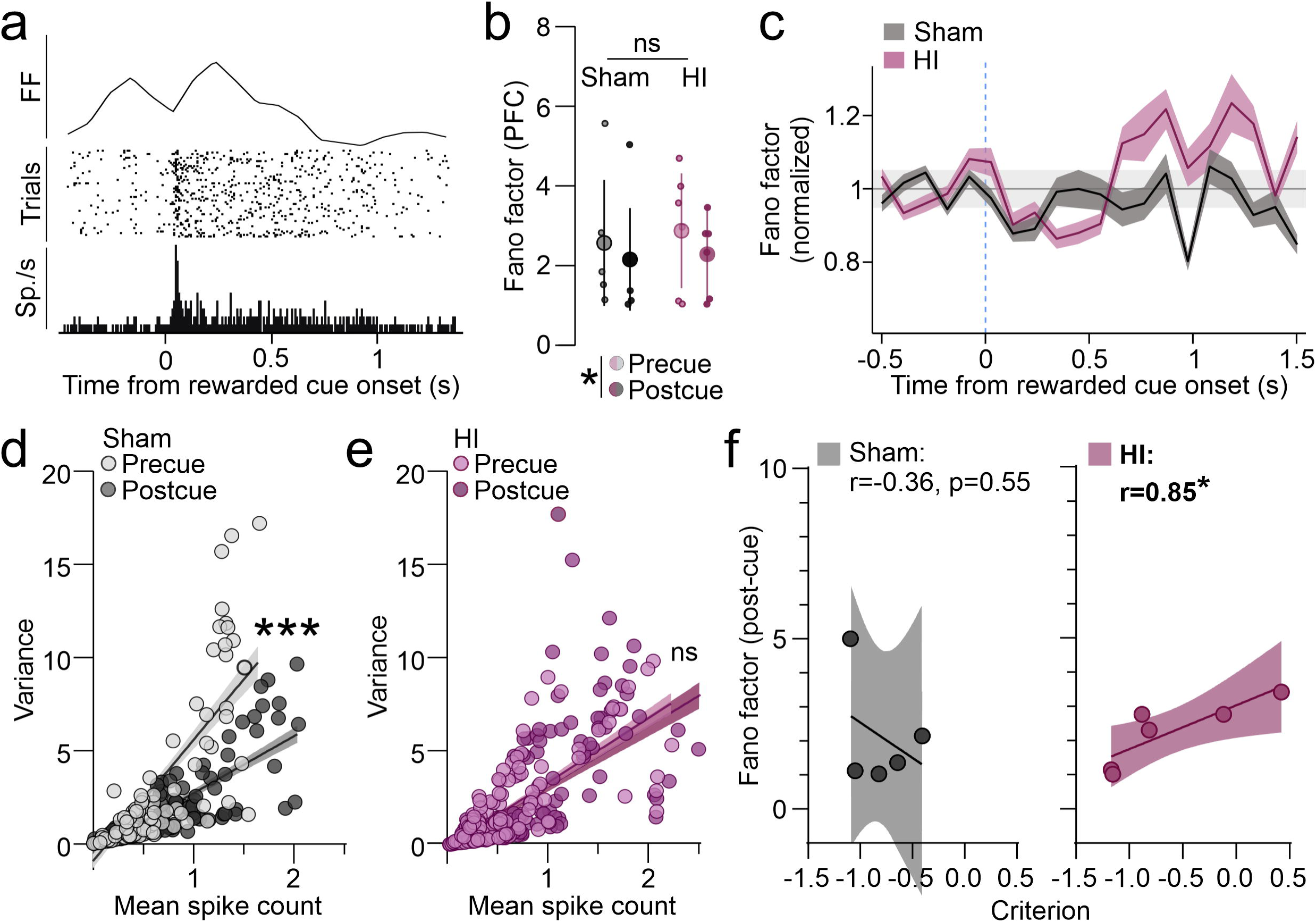
Variability of prefrontal firing has altered temporal dynamics in HI mice and is predictive of behavior. **a)** Representative prefrontal unit and its Fano factor to the rewarded cue (indicated) with a visible quenching after cue onset. **b)** Variability is quenched by the rewarded cue onset in both sham and HI mice (GLM, β_NeonatalHI_=-0.11, p=0.79; β_Cue_=-0.25, p=0.004, β_NeonatalHI×Cue_=-0.04, p=0.55; Hommel’s post-hoc pre-cue vs post-cue sham p=0.08, z=-1.75, HI p=0.009, z=-2.8). **c)** z-scored Fano factor from 0.5 s before the rewarded cue onset and 1.5 s after it, plotted over time, shows higher post-cue variability in HI mice. Grey line indicates baseline±1SD. **d)** Variance and mean spike count have a linear relationship before and after cue onset in sham PFC, with the slope of linear regression significantly lower after the cue onset (linear regression slope_Pre-cue_=6.05, R^2^=0.64; slope_Post-cue_=2.97, R^2^=0.74; Pre-cue vs Post-cue p=0.000). **e)** The slope of the linear regression between variance and mean spike count was unchanged in HI mice (slope_Pre-cue_=4.39, R^2^=0.67; slope_Post-cue_=4.53, R^2^=0.61; Pre-cue vs Post-cue p=0.63). **f)** Post-cue Fano factor values are not correlated with *c* in sham mice (left), unlike in HI mice (Pearson’s r sham=-0.36, p=0.55, N=5 sham mice; r HI=0.85, p=0.033, R^2^=0.72). Data represents individual animals (N=5 sham and 6 HI mice) and 95% CI in b and d-f. c: Data represents mean±SEM of N=183 units from 5 sham and 294 units from 6 HI mice.

Variance and mean spike count of neurons that are used to calculate the Fano factor have a linear relationship in the V1 (Wiener et al., 2001). In the PFC of sham mice, variance and mean spike count have a linear relationship even before the cue onset (Figure 5D; Pearson’s r_Pre-cue_=0.8, p=2.57×10^-41^). Cue presentation increases the mean spike count (GLM, cue p=5.9×10^-4^), reflected in a lower Fano factor after cue onset (Figure 5B) and the slope of the linear regression between variance and mean spike count (Figure 5D; linear regression slope_Pre-cue_=6.05, R^2^=0.64; slope_Post-cue_=2.97, R^2^=0.74; Pre-cue vs Post-cue p=0.000). As Fano factor of HI mice overshoots the baseline between 0.65 and 1.25 s after cue onset (Figure 5C), we hypothesized that the slope of the linear regression between variance and mean spike count would appear unaltered by cue presentation. Indeed, while the linear relationship between variance and mean was preserved in HI mice (Pearson’s r_Pre-cue_=0.82, p=1.6×10^-67^; r_Post-cue_=0.78, p=2.2×10^-57^), the rewarded cue onset did not result in a lower regression slope (Figure 5E; linear regression slope_Pre-cue_=4.39, R^2^=0.67; slope_Post-cue_=4.53, R^2^=0.61; Pre-cue vs Post-cue p=0.63). Despite a lower pre-cue regression slope of variance and mean in HI mice when compared to sham (p=0.000008), neither the mean nor variance values were different between sham and HI mice (GLM; mean spike count p_NeonatalHI_=0.43, variance p_NeonatalHI_=0.89), indicating that neonatal HI selectively impacts their linear relationship. Finally, we tested the association between the post-cue Fano factor and *c* in sham and HI mice. Post-cue Fano factor values in sham mice were not significantly correlated with *c* (Pearson’s r=-0.36, p=0.55, N=5 sham mice). However, Fano factor had a positive association with *c* in HI mice and was a significant predictor of their performance (Pearson’s r=0.85, p=0.033, R^2^=0.72). Altogether, our results demonstrate that neonatal HI injury impairs the temporal dynamics of the Fano factor in the PFC and that the decision bias (*c*) in HI mice has a positive relationship with cue-driven changes in firing rate variability.

## Discussion

In this study, we used an established visual discrimination task in head-fixed mice to uncover the long-term impact of neonatal hypoxic-ischemic (HI) injury on cue processing during goal-directed behavior. We found that adult mice that were subjected to neonatal HI injury have faster reaction times, but that they commit more errors and have a liberal decision bias, despite an overall intact learning trajectory. Electrophysiologically, HI mice show weakened selectivity for task cues in the primary visual cortex (V1) and abnormally high rebound from cue-evoked suppression of neuronal firing rate variability in the prefrontal cortex (PFC), the latter of which was predictive of their decision bias. Our study extends previous investigations of neuronal activity and behavior in mice with neonate HI injury (Du et al., 2026), uncovers an association between abnormal neural variability and behavioral atypicalities in HI mice (Tsikonofilos et al., 2025; Waschke et al., 2021), and highlights the long-term impact of neonatal injury on neuronal activity and goal-directed behaviors.

The Rice-Vannucci model used in our study is a classical paradigm to induce neonatal encephalopathy (Chavez Valdez et al., 2023; Rice et al., 1981; Vannucci & Back, 2022). The structural damage in this model can be highly variable, ranging from mild to severe (Brockmann et al., 2013; Failor et al., 2010). However, our findings of enlarged ventricles and hippocampal damage on the side ipsilateral to the injury in all HI mice, as well as no significant differences in the thickness of the PFC indicate the robustness of our approach and the consistency of the injury. Our findings also suggest that the hippocampus is more vulnerable to the injury than the PFC at this postnatal age (P7-10), in agreement with previous studies (Alexander et al., 2014; McAuliffe et al., 2006). While the PFC had no overt structural damage in the animals used in our study, it is possible that the damage is subtle and can be uncovered through detailed immunohistochemical investigation of synaptic markers, as has previously been published for markers of critical period plasticity (Parmar et al., 2024). Deficits in the structure and density of Parvalbumin interneurons are a frequent finding in rodent models of neonatal HI (Chavez Valdez et al., 2018; Failor et al., 2010; Fowke et al., 2018; Komitova et al., 2013; Parmar et al., 2024; Trnski et al., 2022), and our future studies plan to investigate the fine structure of the PFC and V1 in HI mice, especially in light of their deficits uncovered in this study.

The first novel finding of our study is that neonatal HI injury does not impact discriminability (d’) and learning trajectory, but decision bias (*c*). Given the previous reports of impaired performance, especially in motor tasks or tasks that rely on the animal’s mobility (Du et al., 2026; Marlicz et al., 2024; Ramani et al., 2013), we used a task design that involves headfixation and tracks licks as the motor output to focus solely on measuring cognitive function. In this way, the animals can perform the task even if they underperform in classical motor learning paradigms, such as rotarod. Indeed, the HI mice show intact discriminability, and they reached the learning criterion at the same speed as the sham mice and similar to other control mice from our laboratory (Fariborzi et al., 2026; McCoy et al., 2026). Our results demonstrate that HI injury does not impair associative learning and highlights the need to account for the motor deficits of HI mice when designing studies that test their cognitive abilities (McAuliffe et al., 2006; van der Kooij et al., 2010). Unlike discriminability, other aspects of behavioral performance were altered in HI mice, namely the magnitude of licking responses, the reaction times and decision bias (*c*). While the increased number of licks following the reward, shorter reaction times to both cues and a liberal decision bias may be due to increased motivation in HI mice, the reaction times in HI mice followed a trend similar to sham mice, slowing down as the session progressed and as the mice fatigued, indicating that HI mice are not simply hypermotivated. Their faster responding to both task cues coupled with liberal decision bias and a higher error rate suggests an impulsive-like phenotype, in line with reported associations of perinatal hypoxic injury and attention deficit/hyperactivity disorder (ADHD) (Getahun et al., 2013; Zhu et al., 2016). Future studies can now test the impulsive-like phenotype of HI mice further, such as through start-stop task (SST) and/or delay discounting, both of which can be coupled with recordings of neural activity (Kim et al., 2021; Masuda et al., 2020).

Our recordings of neural activity in this study confirmed previous findings of reduced responsiveness and firing of V1 neurons after the neonatal HI injury (Failor et al., 2010), and extended them to neuronal activity during task performance. Reduced representation of the rewarded cue in regular-spiking neurons, as well as overall reduced selectivity for and responsiveness to task cues suggests persistent sensory processing deficits after neonatal HI injury, in spite of intact feature selectivity (Failor et al., 2010). Interestingly, the responses to the rewarded cue are blunted in both regular and fast-spiking interneurons in HI mice, unlike the responses to the non-rewarded cue. A previous study demonstrated that increased excitability in the V1 during cue presentation results in increased Hits and False Alarms, as well as a liberal decision bias (Jin & Glickfeld, 2019). Our findings of reduced cue-evoked firing, increased False Alarm rate and liberal decision bias in HI mice seemingly contradict this study. However, our results from sham mice support that neural activity in the V1 correlates with decision bias, specifically the cue-evoked firing rate of putative fast-spiking neurons, as well as the ratio of visually-evoked potential (VEP) responses to task cues, the latter of which is a novel finding and confirms the utility of VEPs in the diagnosis of neurodevelopmental outcomes (Cainelli et al., 2018). Our results hence indicate that the relationships between neural activity and behavior found in the neurotypical brain do not apply to neurodevelopmental conditions. Indeed, a previous study reported a lack of correlation between neuronal activity in the hippocampus and rotarod performance in HI mice, in contrast to sham mice (Du et al., 2026). Altogether, our study underscores the need for detailed studies of neural activity in animal models of neurodevelopmental conditions, especially during cognitive tasks.

Task-related neuronal activity in the V1 seemed more impacted by neonatal HI injury than the activity in the PFC, a brain area thought to mediate the goal-directed behaviors (Du et al., 2026; Failor et al., 2010). Representation of and selectivity for task cues appeared almost intact in the prefrontal neurons, apart from fast-spiking interneurons, whose pre-cue firing was reduced. This finding is in line with deficits in Parvalbumin interneurons after neonatal HI injury and is common to both V1 and PFC in our study, highlighting their vulnerability to neonatal injury (Failor et al., 2010). However, HI mice had altered temporal dynamics of neural firing rate variability in the PFC, measured as the Fano factor (Jin & Glickfeld, 2019). While the cue onset quenched variability in firing in both groups of mice, firing variability in HI mice fluctuated significantly more than in sham mice and overshot the baseline values by almost 25%. Interestingly, the decision bias of HI mice was highly correlated with their Fano factor, unlike in sham mice, indicating that subtle impairments of prefrontal activity contribute to their behavioral phenotype. To the best of our knowledge, our findings provide a first relationship between the Fano factor and behavior and highlight its utility in studying neuropsychiatric conditions (Tsikonofilos et al., 2025; Waschke et al., 2021). Future studies will undoubtedly address the mechanisms that regulate the Fano factor dynamics in different brain areas and its association with behavioral and cognitive deficits.

Due to the homogeneity of injury in our model, our study was unable to correlate the heterogeneity of HI injury with behavior. Further, the late postnatal timing of injury in our model may not model the HI injury in preterm or early term neonates. Despite these limitations, our study identified persistent behavioral atypicalities and impairments in the processing of visual cues in the V1 of mice after neonatal HI injury. Lasting impact of neonatal HI injury underscores the need for detailed studies of developmental trajectories after perinatal and neonatal injury, especially considering differential vulnerability of brain areas. Our study further identified an association between neural variability in the PFC and behavioral decision bias in HI mice, informing future studies in human populations focused on identifying biomarkers of brain injury (Tsikonofilos et al., 2025).

## Methods

### Mice

Mice were maintained on C57BL/6 background (Charles River Laboratory, Wilmington, MA) on reverse 12:12 light:dark cycle (lights off 11 AM-11 PM), with food and water *ad libitum,* except during visual discrimination training (see below). HI injury was done on postnatal day (P) 10 as described previously using the Vannucci model adapted to mice (permanent unilateral left carotid artery ligation, plus 60 minutes of hypoxia at FiO2 = 0.08) (J. Burnsed et al., 2019; J. C. Burnsed et al., 2015; Ditelberg et al., 1996). For carotid ligation surgery, mice were anesthetized using inhaled isoflurane (3% for induction, followed by 1% maintenance). Pups were returned to their dams for a 1h rest period between surgical ligation and hypoxia. Both male and female mice were used in the study and the experimental groups were balanced in terms of sex. Animals were treated in accordance with the University of Virginia Institutional Animal Care and Use Committee guidelines.

### Headpost implantation surgeries

Animals were anesthetized with isoflurane in oxygen (2-2.5% induction, 1–1.5% maintenance), warmed with a heating pad at 38°C and given subcutaneous injections of Rimadyl (5 mg/kg) and 0.25% Bupivacaine (beneath their scalp). Eyes were covered with Puralube (Decra, Northwich, UK). Scalp and fascia from Bregma to behind lambda were removed, and the skull was cleaned, dried and covered with a thin layer of Scotchbond adhesive (3M, Maplewood, MN). Skin edges were sealed with VetBond (3M). The head plate (stainless steel, SendCutSend, Reno, NV) was attached with dental cement (RelyX Ultimate, 3M). After the cement was cured, the well of the head plate was filled with silicone elastomer (Reynold Advanced Materials, Brighton, MA) to protect the skull.

Mice were given Rimadyl dissolved in hydrogel (1% food-grade agar in distilled water) during the recovery (Ingrao et al., 2013). Animals were group-housed after the implantation and monitored daily for signs of shock or infection. On the day of electrophysiological recording, the animals were anesthetized as above and craniotomies (∼0.5 mm in diameter) were made above V1, PFC and cerebellum with 18G needles. The brain surface was covered in 2-3% low melting point agarose (Promega, Madison, WI) in sterile saline and then capped with silicone elastomer. Animals were allowed to recover for 1.5-2 h before the recording.

### Electrophysiology

For the recording sessions, mice were placed in the head-plate holder above an aluminum mesh treadmill and allowed to habituate for 5-10 minutes. The silicone plug was removed, the reference (insulated silver wire electrode; A-M Systems, Carlsborg, WA) was placed in the cerebellum and the well was covered with warm sterile saline. A multisite electrode spanning all cortical layers (A1×16-5mm-50-177-A16; Neuronexus Technologies, Ann Arbor, MI) was coated with DiI (Invitrogen) to allow post hoc insertion site verification and then inserted in the V1 through the craniotomy. For PFC, a 4×4 shank probe (A4×4-3mm-50-125-177-A16) was used for the recordings, also coated with DiI. The electrodes were slowly (5-10 µm/s) lowered to the appropriate depth: 800 µm for the PFC and until the uppermost recording site had entered the brain for the V1 and allowed to settle for 15-30 minutes. The well was filled with 3% agarose prior to the recordings to stabilize the electrode and the whole region was kept moist with surgical gelfoam soaked in sterile saline (Pfizer, MA). The signals from the recording probes were fed into a 16-channel amplifier (Model 3500; A-M Systems, Sequim, WA) and amplified 200x, before being sampled at 25 kHz using Spike2 and Power 1401-3 data acquisition unit (CED). After the recording ended, the electrodes were slowly retracted, and the well was cleaned and protected with silicone elastomer.

### Visual stimuli

For visual stimuli, blank screen was generated with MATLAB (MathWorks, Natick, MA) using the PsychToolBox extension (Brainard, 1997) and presented on a gamma corrected 27” LCD. The screen was centered 20-25 cm from the mouse’s right eye, covering ∼80° of visual space. For visual discrimination training, visual stimuli (120° and 60° gratings) were presented at 0.15 cpd and 100% contrast in a random sequence for 3 seconds, followed by a 15 second long interstimulus interval (blank screen). Presentation of 120° triggered the delivery of 10 µl of water through a syringe pump (New Era Pump Systems, Farmingdale, NY), while the presentation of 60° had no consequence.

### Behavior

4-7 days after the headpost implantation, mice were gradually water-restricted (from 3 ml of water/day to 1 ml of water/day) for 5-7 days. Food was available *ad libitum*. Mice were weighed daily and their weight was maintained at 85% of initial weight to prevent dehydration. During water restriction, mice were gradually habituated to handling by the experimenters, the treadmill, and the water delivery spout during daily habituation sessions (Guo et al., 2014). After water restriction, the visual discrimination training began, where mice were headfixed above the treadmill and a stainless-steel gavage needle (18G) was positioned near the mouse’s mouth for water delivery. The pump was controlled by the Power1401-3 data acquisition interface (CED) for lick detection (Hayar et al., 2006). Mice were trained in 2 daily sessions consisting of 50 120° and 60° presentations each, for a total of 200 trials per day. Licks of the spout during the 10 s after the onset of stimulus presentation were quantified according to the signal detection theory as Hits (the fraction of trials in which the mouse licked to the 120° orientation), Misses (the fraction of trials in which the mouse failed to lick to the 120° orientation), False Alarms (the fraction of trials in which the mouse licked to the 60° orientation) and Correct Rejections (the fraction of trials in which the mouse failed to lick to the 60° orientation), where the performance was measured as discriminability (d’)=z(H) - z(FA). Decision bias *c* was calculated as-0.5×[z(H)+z(FA)]. Mice were trained until they achieved a d’ of 2 or above for 3 consecutive sessions.

Animals were monitored for signs of distress for the duration of the experiment. Animals that developed corneal damage, cataracts, and symptoms that had the potential to impact their behavior or physiology (such as immobility and other signs of pain, sudden weight loss or impeded weight gain) were excluded from the study. All 18 sham mice and 10 HI mice underwent craniotomies for electrophysiology and/or the eletrophysiological recordings, but only 8 sham and 8 HI mice had satisfied the criteria for inclusion in recordings and analysis, namely 1. confirmed recording locations in the V1 and the PFC, 2. absence of motion artifact that interfered with data analysis, 3. absence of electrical noise, 4. absence of profuse bleeding from the superior saggital sinus after the opening of the craniotomy, 5. absence of pulsation artifact from the saggital sinus during the recording, and 6. satisfactory performance during the recording session (for example, no spout grabbing or overt grooming as it precludes the lick analysis).

### Histology and imaging

After the last training session or electrophysiological recording, mice were anesthetized with a mixture of ketamine and xylazine and transcranially perfused with warm 0.1 M phosphate buffer, followed by warm 4% paraformaldehyde (Electron Microscopy Sciences, Hatfield, PA). Brains were postfixed 1 hr at room temperature, followed by overnight fixation at 4°C. Brains were sectioned into 100 µm sections using a vibratome and stored in 1x phosphate buffered saline (PBS) and 0.01% sodium azide. For quantification of PFC thickness, every 5^th^ section until Bregma from each mouse was taken and incubated in NeuroTrace 435/455 (Thermo Fisher Scientific, Waltham, MA) 1:1000 for 2 hours at room temperature in 1x PBS. The left hemisphere was labeled with a longitudinal cut on the ventral side made with an 18G needle dipped in DiO (Thermo Fisher). Before imaging, the sections were rinsed in distilled water, mounted on glass slides, briefly dried, and coverslipped with Aquamount (Polysciences, Warrington, PA). Images were acquired using Leica Stellaris 8 at 2048×2048 resolution using 10x HC PLAN FLUOTAR air (NA=0.3 for tiling) and stitched using LASX software. PFC thickness was measured manually using ImageJ by measuring the distance from the cortical surface to the edge of the corpus callosum, while ensuring that the line was drawn under a 90° angle from the cortical surface to ensure consistency. PFC thickness was measured from posterior PFC [areas Cg1 and Cg2 from (Paxinos & Franklin, 2019)]. All measurements from a single mouse were averaged per hemisphere before statistical comparison.

### Electrophysiological and behavioral data analysis

LFPs were analyzed using waveform averaging from raw, unfiltered traces using Spike2 (CED). Spikes were isolated from LFP recordings using template matching in Spike2. Briefly, the recordings were band-pass filtered (0.7-7 kHz) and smoothed (1 ms). Threshold for detection was set at 4 SDs of the mean baseline (blank screen) and a window of 0.9-1 ms. Isolated units were clustered using waveform properties (amplitude, spike half-width and slope of repolarization), and the clusters were checked manually for quality in addition to confirming there were no spikes during the refractory period (2 ms) using interval histograms. Units with refractory period violations or units with large variations in amplitude (>10%) were excluded from analysis. Out of 8 sham and 8 HI mice that were included in the electrophysiology analysis, units were isolated from 5 sham and 6 HI mice, where 1 sham mouse had no identifiable fast-spiking waveforms. Selectivity indices (SIs) were calculated from 10 ms peristimulus time histogram (PSTHs) as (R_cue_-R_baseline_)/(R_cue_+R_baseline_), where R_cue_ is the firing rate during the initial 1-1.5 s presentation of 120° and 60° stimuli. Cue-evoked firing rates were calculated as baseline firing (0.5 seconds prior to cue onset) subtracted from the the peak firing after the cue (within a 1.5 s window). Fano factor was calculated as variance/mean spike count of 100 ms bins of trial-to-trial firing (0.5 s prior and 1.5 s following the cue onset). Stationary and moving stages were not analyzed separately as animals were typically walking in short bouts during the recordings. Recordings where large motion artifacts were present when the animal was simply balancing on the treadmill or grooming were excluded from analysis, along with recordings from locations outside of the PFC or V1 (as determined through post-hoc mapping of insertion sites with DiI).

### Quantification and statistical analysis

All analyses were performed with the researchers blind to the condition. Statistical analyses were performed in GraphPad Prism 9.0 (GraphPad Inc., La Jolla, USA), R and JASP (JASP Team 2026). N refers to the number of animals or single units, as indicated in figure legends or in text. Behavioral data was analyzed using LMMs with group as a fixed factor and animal as a random factor. For SI analysis, units significantly modulated by cues were determined through comparison of pre-cue and post-cue firing using Wilcoxon signed rank test. The proportions of significantly modulated units (positively and negatively) were determined and reported per animal, and PSTHs and SIs represent significantly modulated units. PSTHs were compared using Mann-Whitney t-test test of area under the curve (AUC) calculated from 0.5 s precue and 1.5 s postcue activity (10 ms bins). All of the unit data failed the normality tests, so GLMs were used for their analysis (Identity or Log link function) with group and cue as fixed factors and animal as random factor. p-values of contrasts were adjusted using Hommel correction. For correlation and linear regression, SI, cue-evoked and Fano factor values from all neurons per animal were averaged and used as predictors of *c* as described in text and figure legends. CUMSUM was conducted in R on a per-neuron baseline-normalized Fano factor values. PELT changepoint detection was conducted in R using *changepoint* package (Killick & Eckley, 2014) on z-scored Fano factor values. Data are reported as mean ± 95% confidence interval or SEM (standard error of mean), as indicated in text or figure legends, where N represents the number of animals or the number of single units (as indicated).

## Data Availability

All raw data can be obtained by contacting the corresponding author.

## Code Availability

All analysis codes used in this study are available online (CED website) under https://ced.co.uk/downloads/scriptspkexpr

## Author Contributions

**S.N.** Investigation, Formal Analysis. **M.K.** Investigation. **H.M.M.** Investigation. **N.K.S.** Formal Analysis. **J.C.B.** Methodology, Resources, Writing-Review & Editing. **A.R.** Conceptualization, Methodology, Formal Analysis, Writing-Original Draft, Visualization, Supervision, Funding Acquisition.

## Acknowledgements

This study was supported by the Department of Pediatrics at the University of Virginia to J.C.B, and the Department of Psychology at the University of Virginia, Owens Family Foundation and R01MH140184 to A.R. and The authors would like to thank the Program in Fundamental Neuroscience for generous access to their confocal microscope.

---

**Supplementary Figure 1. Motivation during task is similar between sham and HI mice.** Reaction times show a similar increase as the session progresses in both sham and HI mice (LMM, β_trial_=-0.038, p=0.037, N=18 sham and 10 HI mice).

**Supplementary Figure 2. SI values are not correlated with decision bias.** SI values of RS **(a)** or FS **(b)** neurons are not correlated with c in either sham or HI mice. RS Pearson’s r: sham_Reward_=0.89, p=0.1; sham_NoReward_=-0.88, p=0.12; HI_Reward_=0.48, p=0.32; HI_NoReward_=0.6, p=0.2. FS Pearson’s r: sham_Reward_=0.29, p=0.7; sham_NoReward_=--0.91, p=0.09; HI_Reward_=-0.13, p=0.8; HI_NoReward_=0.14, p=0.79. N=4/5 sham and 6 HI mice, as indicated in main text.

Data represent individual points and 95% CI of linear regression.

**Supplementary Figure 3. PFC thickness is intact in HI mice despite hippocampal damage. a)** Representative coronal PFC sections of sham and HI mice ipsilateral to the injury side (left hemisphere) depicting where the PFC thickness was measured. **b)** PFC thickness is not impacted by HI injury (mean thickness sham=958.44±38.73 µm, HI=934.39±52.45 µm; N=12 sham and 10 HI mice, student’s t-test p=0.71, t=0.38, df=20). **c)** Sections from HI mice from anterior and medial hippocampus depicting enlarged lateral ventricle (white arrowhead).

