## Supplementary figures and images for "Persistent sensory processing and behavioral atypicalities in a mouse model of neonatal encephalopathy"

### Supplementary Figure 1

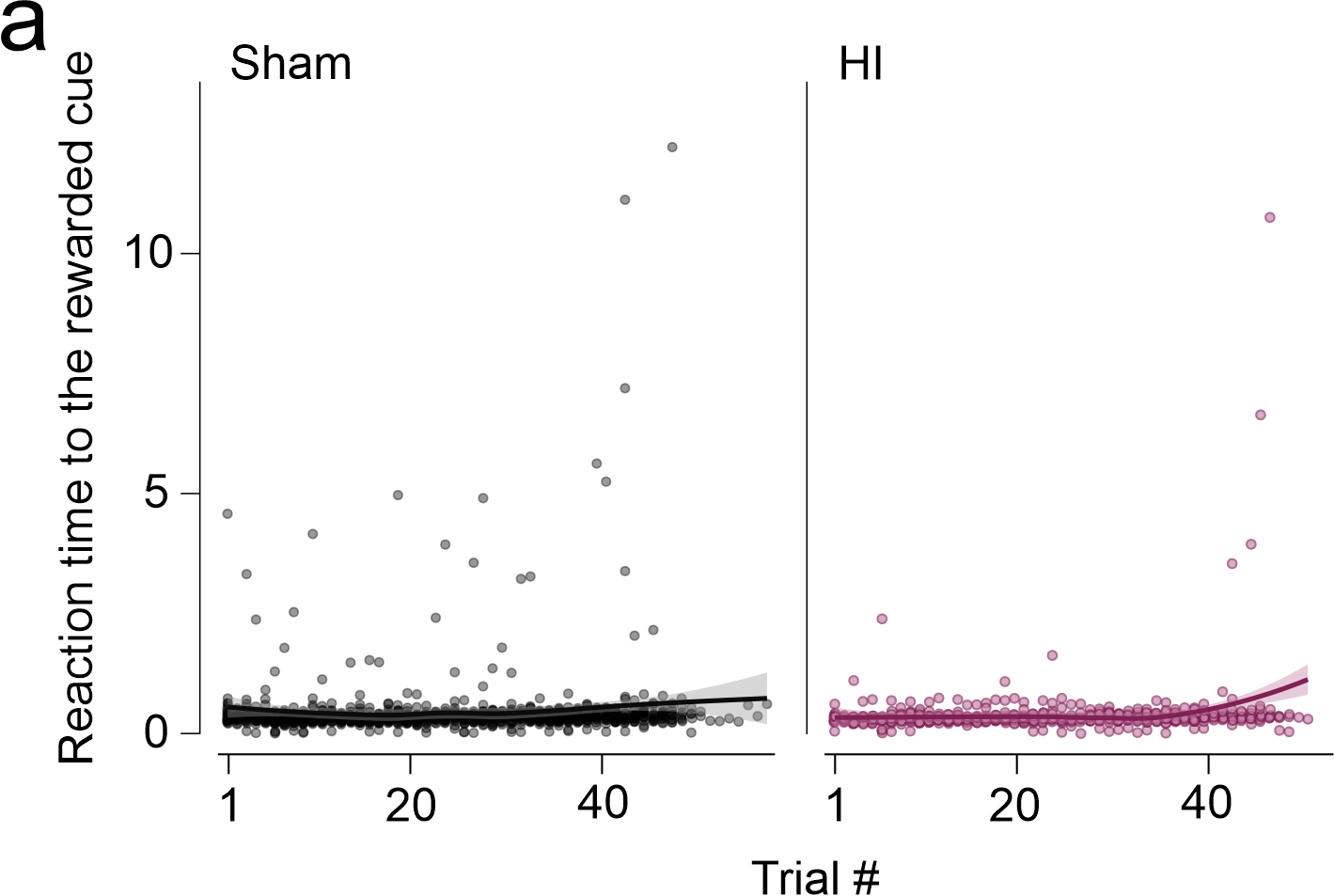

### Supplementary Figure 2

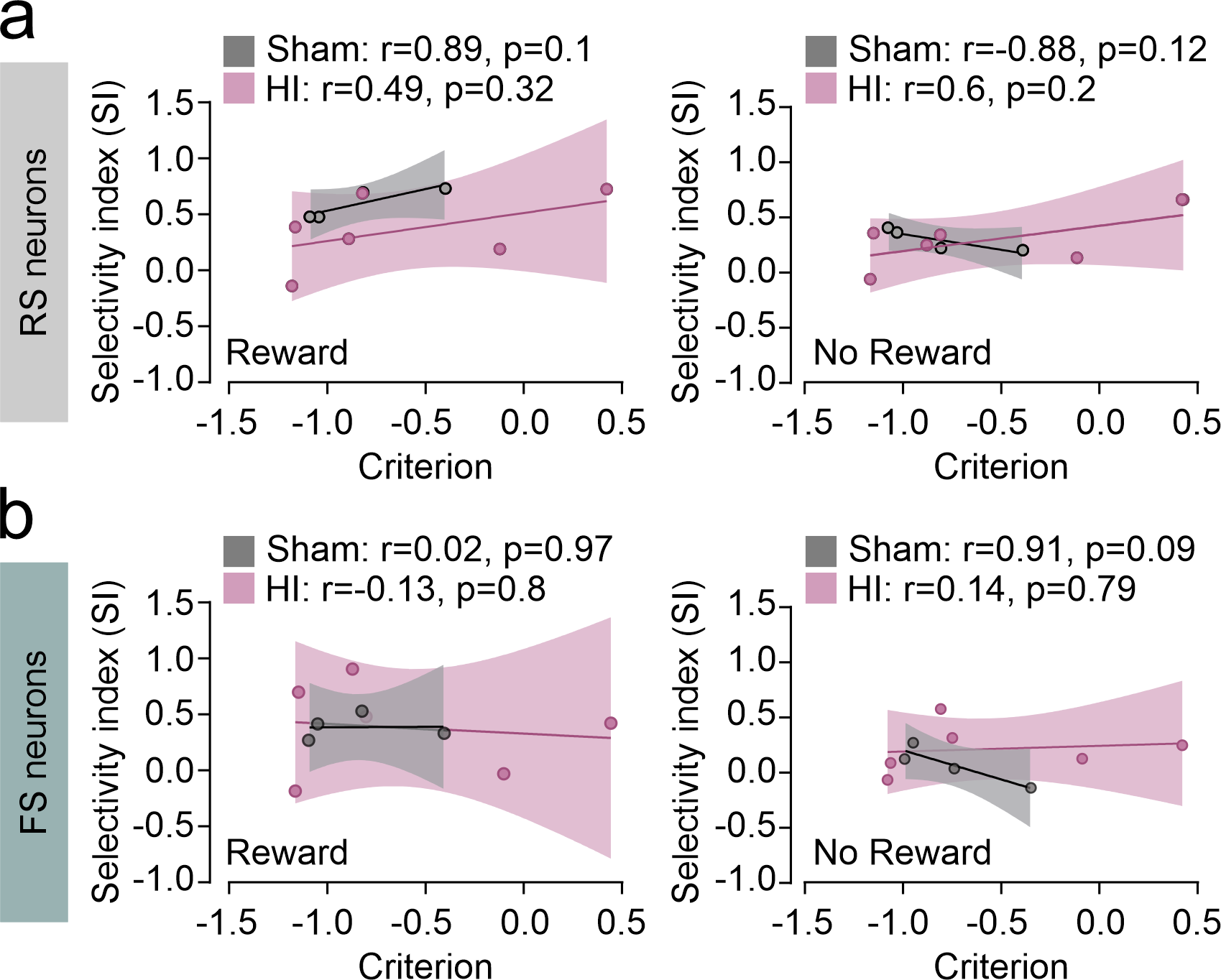

### Supplementary Figure 3

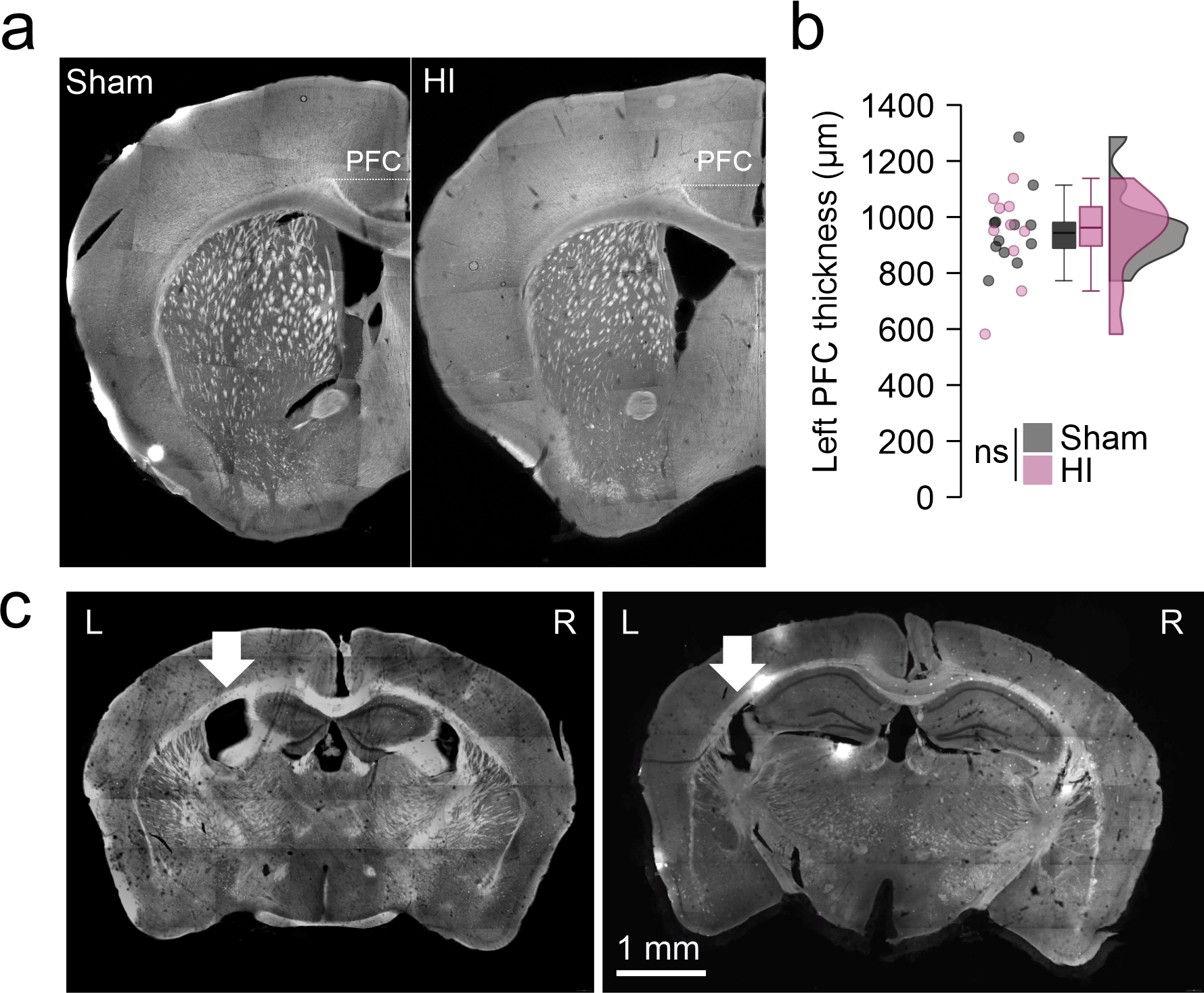
